# A phenotypic small-molecule screen identifies halogenated salicylanilides as inhibitors of fungal morphogenesis, biofilm formation and host cell invasion

**DOI:** 10.1101/361139

**Authors:** Carlos Garcia, Anaïs Burgain, Julien Chaillot, Émilie Pic, Inès Khemiri, Adnane Sellam

## Abstract

A poorly exploited paradigm in the antimicrobial therapy field is to target virulence traits for drug development. In contrast to target-focused approaches, antivirulence phenotypic screens enable identification of bioactive molecules that induce a desirable biological readout without making *a priori* assumption about the cellular target. Here, we screened a chemical library of 678 small molecules against the invasive hyphal growth of the human opportunistic yeast *Candida albicans*. We found that a halogenated salicylanilide (N1-(3,5-dichlorophenyl)-5-chloro-2-hydroxybenzamide) and one of its analog, Niclosamide, an FDA-approved anthelmintic in humans, exhibited both antifilamentation and antibiofilm activities against *C. albicans* and the multi-resistant yeast *C. auris*. The antivirulence activity of halogenated salicylanilides were also expanded to *C. albicans* resistant strains with different resistance mechanisms. We also found that Niclosamide protected the intestinal epithelial cells against invasion by *C. albicans*. Transcriptional profiling of *C. albicans* challenged with Niclosamide exhibited a signature that is characteristic of the mitochondria-to-nucleus retrograde response. Our chemogenomic analysis showed that halogenated salicylanilides compromise the potential-dependant mitochondrial protein translocon machinery. Given the fact that the safety of Niclosamide is well established in humans, this molecule could represent the first clinically approved antivirulence agent against a pathogenic fungus.

## Introduction

*Candida albicans* is an ascomycete fungus that is an important commensal and opportunistic pathogen in humans. Systemic infections resulting primarily from this yeast and the filamentous fungus, *Aspergillus fumigatus*, are associated with mortality rates of 50% or greater despite current therapies ^1-3^. Therapeutic options are limited to treatment with mainly three antifungal classes, namely polyenes, azoles and echinocandins ^4^. These compounds target the specific fungal biological process of ergosterol metabolism (azoles and polyenes) and cell wall β-1,3-glucan synthesis (echinocandins). However, these drugs have serious side effects such as nephrotoxicity and/or create complications such as resistance and interactions with other commonly prescribed drugs. There are currently a limited number of novel antifungal molecules on the drug discovery pipelines and most of them target the same cellular processes as azoles and echinocandins ^5-7^. Thus, these molecules will most likely face the same limitations in term of resistance and toxicity. These considerations highlight the urgent need to identify novel targets and strategies for antifungal therapies

The pathogenicity of *C. albicans* is mediated by different factors such as invasive filamentation, biofilm formation and the ability to escape the immune system ^4^. A relatively new but poorly exploited paradigm in the antimicrobial therapy field is to target virulence traits for drug development ^8,9^. In contrast to target-focused approaches, phenotypic screens enable identification of bioactive molecules that induce a desirable biological readout without making *a priori* assumption about the cellular target. Additionally, compounds identified by phenotypic screens are by definition cell permeable and engage their target with sufficient affinity. Taking into consideration the commensal lifestyle of *C. albicans*, suppressing its growth in different niches inside the host will perturb the microbial flora equilibrium and lead to other opportunistic infections ^10^. Specific inhibition of fungal virulence determinants without affecting commensal growth represents thus an attractive approach.

*C. albicans* is a polymorphic fungus that are able to reversibly shift to different morphologies including yeast, pseudohyphae and true hyphae forms. Hyphae are long tubular cells that are associated with the invasion of organs and tissues of the human host during Candidemia episodes.

In addition to ensuring a physical force required to penetrate host cells, the hyphae state is characterized by an enhanced adhesiveness and allow to the fungal cells to escape the from phagocytes and to conquer niches where nutrient conditions are not limiting ^11^. Furthermore, even if hyphae are not essential for the formation of *C. albicans* biofilms, they are determinant for the compression strength and the resistance to mechanical dislocation of this highly resistant growth lifestyle ^12^. Thus, antivirulence therapy based on molecules that inhibit hyphae formation could have substantial benefits on managing fungal infections. Moreover, antivirulence agents might provide an alternative strategy to circumvent antifungal resistance by disarming fungal resistant pathogens from their virulence factors ^13^.

Proof of principle for this concept has been demonstrated in different investigations ^14-19^. Fazly *et al*. characterized a small molecule called filastatin that inhibits *C. albicans* filamentation, adhesion and virulence ^15^, yet, the molecular mechanism by which this molecule acts remain elusive. Recently, screening of small molecules that are being clinically tested for different pathologies identified two antifilamentation compounds that also perturbed endocytosis ^16^. However, these compounds were also found to inhibit the commensal yeast growth at the effective antifilamentation concentrations.

In this study, we screened a chemical library of 678 small molecules that were preselected for bioactivity in the budding yeast *S. cerevisiae* and have well-balanced hydrophilic-lipophilic properties allowing the crossing of both the hydrophilic fungal cell wall and the lipophilic membrane. We found that N1-(3,5-dichlorophenyl)-5-chloro-2-hydroxybenzamide (TCSA: <u>T</u>ri-<u>C</u>hloro-<u>S</u>alicy<u>a</u>nilide, for simplified nomenclature), a halogenated salicylanilide, was a potent antifilamentation molecule and inhibited also biofilm formation of both *C. albicans* and the multi-resistant yeast *C. auris*. The TCSA analog, Niclosamide, that is an FDA-approved anthelmintic agent, exhibited a similar anti-filamentation and anti-biofilm activities and conferred a significant protective activity to intestinal epithelial cells against fungal invasion. Both genome-wide transcriptional profiling and chemical-genetic approach were used to gain insight into the mechanism of action associated with the antivirulence activity of the halogenated salicylanilides (HSA). We found that HSA-mediated antivirulence activity is likely related to the perturbation of the fungal mitochondrial protein import machinery.

## Results

### Phenotypic screening of yeast-bioactive small molecules for inhibition of hyphal growth

To identify small molecules with antivirulence properties the yeast bioactive small molecule library ^20^ was screened for compounds that inhibit hyphal formation (Fig. 1A). Compounds of this chemical library were preselected for bioactivity in *S. cerevisiae* and have well-balanced hydrophilic-lipophilic properties allowing the crossing of both the hydrophilic fungal cell wall and the lipophilic membrane. *C. albicans* cells were grown in 96-well under hyphae-promoting conditions (SC-10% fetal bovine serum (FBS)) for 3h at 37 °C in the presence of the bioactive compounds. The ability of each compounds to inhibit *C. albicans* SC5314 strain morphogenesis was assessed by imaging each well using the high-content microscope Cytation 5. This primary screen identified 93 molecules with anti-filamentation activity at 100µM (70-80% of *C. albicans* cells with no germ tubes). A total of 50 promiscuous molecules with chemical structures similar to pan assay interference compounds (PAINS) associated with antifungal activity ^21^ were removed from the hit list of the primary screen. Inhibition of filamentation could be associated with the fact that small molecules are very toxic which might systematically alter all developmental process of a fungal cell such as morphogenesis. To rule out cytotoxic molecules that impaired growth *per se*, the 43 retained hits were assayed for their ability to inhibit growth of yeast cells growing at 30°C. Through this process, a subset of five molecules were selected which were then assayed for their potency using a quantitative dose-response filamentation inhibition assay. Among the five hits, we found two molecules that had a significant effect on filamentous growth at pharmacological concentrations of 1-10µM (data not shown). The effect of this compounds on *C. albicans* filamentation were confirmed using a new batch of powders from different suppliers (Fig. 1B-C).

**Figure 1.**
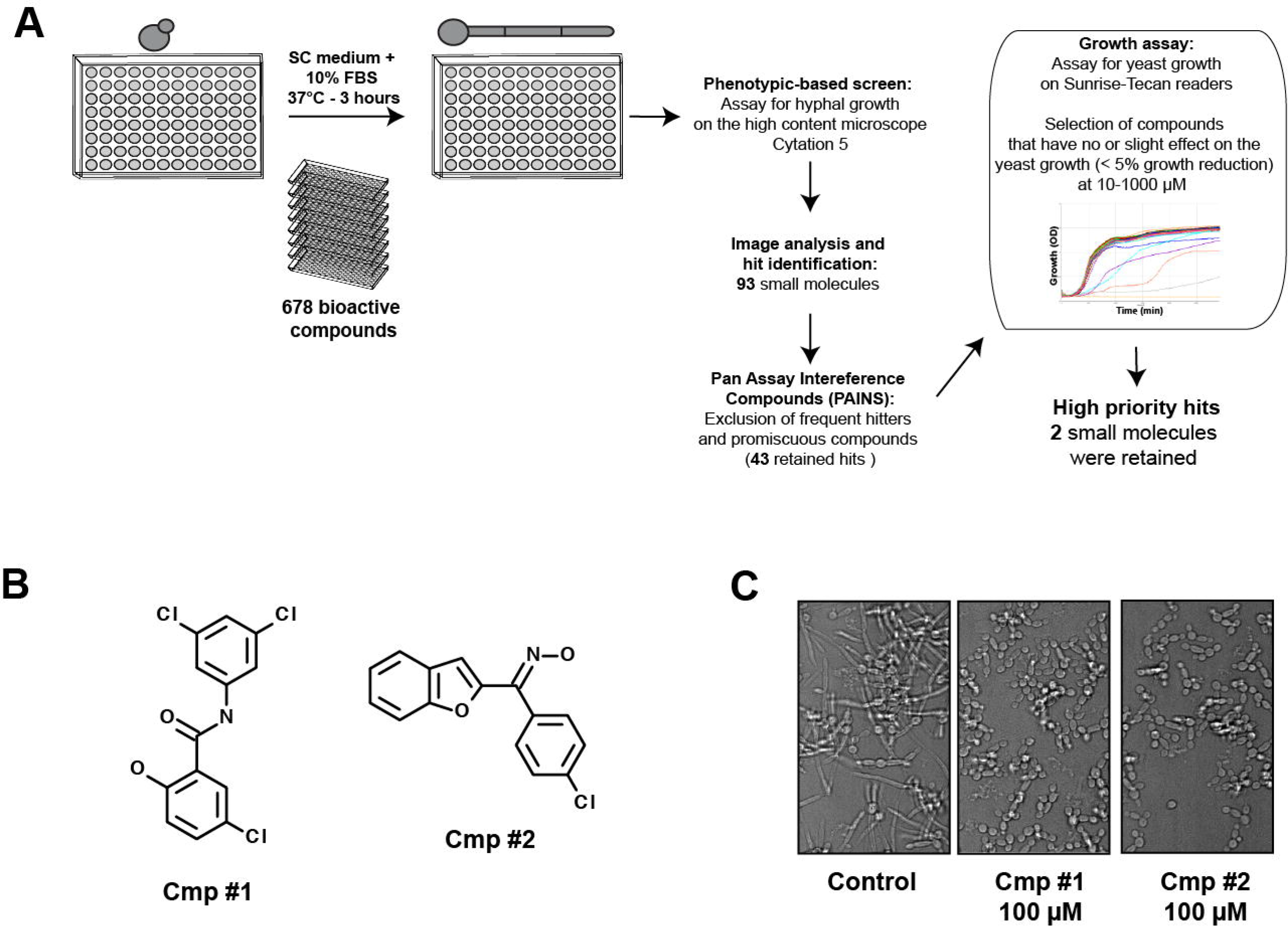
Hyphal growth inhibition assay in *C. albicans*. (**A**) Workflow of small-molecule high-throughput screen for hyphal growth inhibition of *C. albicans* SC5314 strain. (**B**) Chemical structure of the antifilamentation compound 1 (Cmp #1; N1-(3,5-dichlorophenyl)-5-chloro-2-hydroxybenzamide) and (Cmp #2; 3-Chloro-aryl(benzofuran-2-yl) ketoxime). (**C**) Antifilamentation activity of the Cmp #1 and Cmp #2 at 100 µM. *C. albicans* SC5314 strain was grown under hyphae-promoting conditions (SC medium supplemented with 10% FBS) for three hours at 37°C.

Compound #1 (N1-(3,5-dichlorophenyl)-5-chloro-2-hydroxybenzamide or TCSA) is a halogenated form of salicylanilide scaffold. Salicylanilides are widely used as antiparasitic veterinary drugs against helminths and ectoparasites ^22^. The salicylanilide Niclosamide (5-chloro-salicyl-(2-chloro-4-nitro) anilide) was approved by the FDA to treat tape worm infection in humans ^23^ and has been recently shown to have anti-cancer and anti-diabetic activities ^24,25^. Recently, HSA including Niclosamide and Oxyclozanide were repurposed as alternative antibiotic therapy against clinical resistant isolates of *Staphylococcus aureus* ^26^. The compound #2 (3-Chloro-aryl(benzofuran-2-yl) ketoxime) had chemical features with previously described biological activities. This compound had both oxime and benzofuran residues that are associated with the antifungal activity of many molecules ^27-32^. Derivatives of compound #2 were also shown to be active against *C. albicans* by targeting the N-myristoyltransferase, Nmt1 ^33,34^. While in our conditions this molecule had a slight effect on the growth of the yeast form of *C. albicans* (10% of inhibition at 1mM), other studies using different growth medium (RPMI) had shown a strong growth reduction at 15µM ^31^. Based on this, we decided to focus our downstream investigation on compound #1.

### TCSA and Niclosamide inhibit *C. albicans* filamentation without affecting the commensal yeast growth

Commercially available HSA analogs 3-12 were purchased and tested on both growth and filamentation (Fig. 2A). With the exception of Niclosamide (compound #6), all tested salicylanilide derivatives impaired the growth of *C. albicans* at concentration of 20µM (>50% inhibition of the growth as compared to the control) (Fig. 2B). Interestingly, the salicylanilide scaffold alone had a strong effect on the fungal growth, suggesting that halogenation potentiates the antifilamentation rather than yeast antigrowth effect of TCSA and Niclosamide. Both TCSA and Niclosamide have no significant impact on the growth of the yeast form (Fig. 3A-B). At 50µM, TCSA inhibited hyphae elongation by 90%, as compared to the control, while Niclosamide abolished completely *C. albicans* filamentation (Fig. 3C-D).

**Figure 2.**
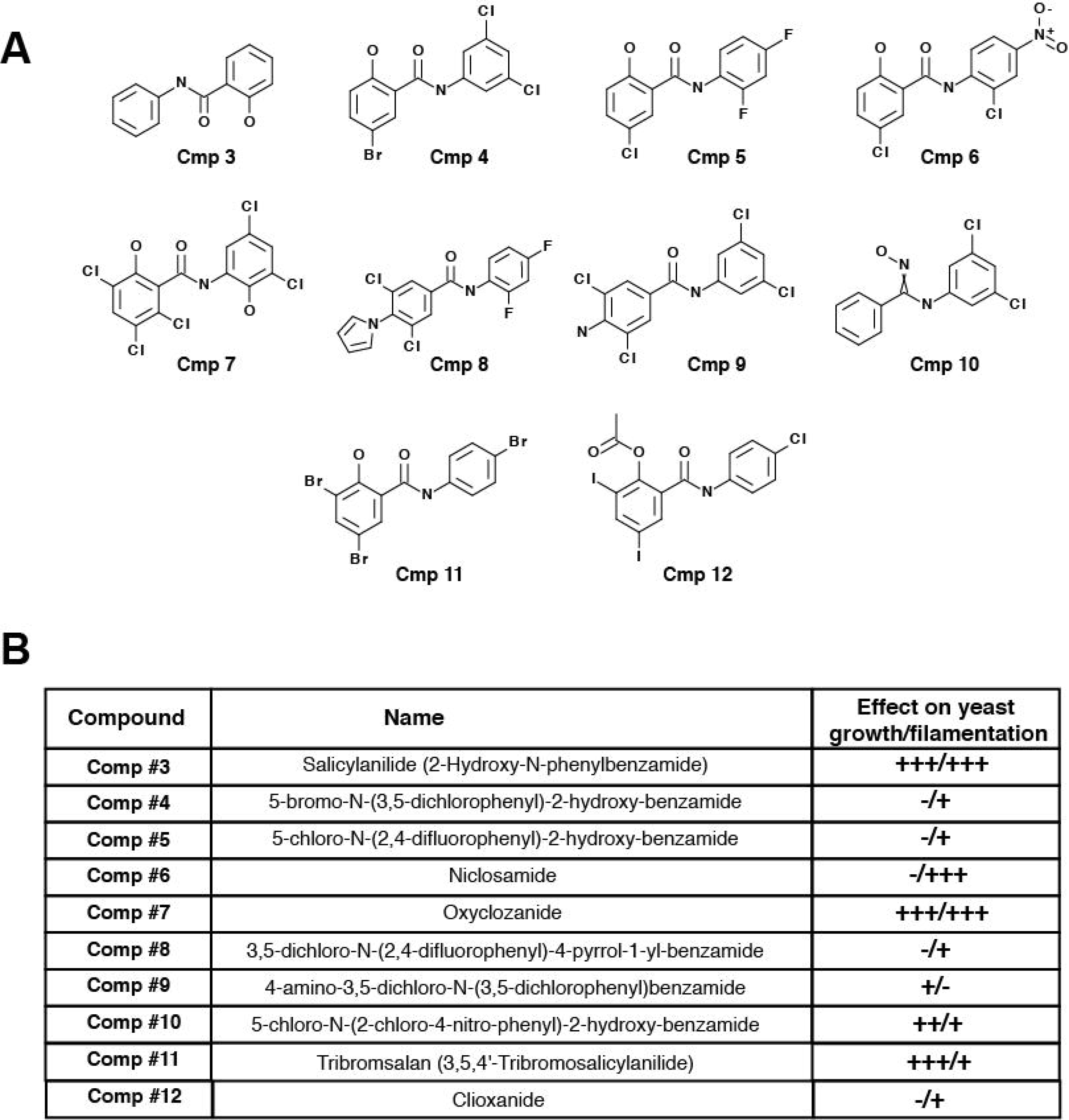
Effect of halogenated salicylanilide analogs on *C. albicans* yeast and hyphal growths. (**A**) Chemical structures of different commercially available salicylanilides. (**B**) Effect of TCSA analogs on both yeast and hyphal growths of *C. albicans*. To assess the effect of each compound on yeast growth at concentrations ranging from 1 to 100µM, *C. albicans* SC5315 strain was grown on SC medium at 30°C for 24 hours. Hyphae-promoting conditions were obtained by incubating *C. albicans* cells on SC medium supplemented with 10 % FBS at 37°C for 3 hours and salicylanilides (1-100µM). (+++), (++) and (+) indicate strong, medium and low inhibition, respectively. Absence of inhibition was indicated by (-).

Since the yeast-to-hyphae transition is triggered by different environmental cues, the antifilamentation effect of TCSA and Niclosamide was also tested and confirmed using other hyphae-promoting media. In RPMI medium, both compounds were effective at concentrations similar to those in SC supplemented with 10% FBS (Fig. 3E). Interestingly, in Spider, Nacetylglucosamine (GlcNAc) and Lee’s media the antifilamentation activity of TCSA and Niclosamide were enhanced and a complete abolition of the *C. albicans* hyphal growth were perceived at concentration ranging from 0.5 to 5µM (Fig. 3F-I). Both TCSA and Niclosamide exhibited antifilamentation activity on other non-*albicans* Candida species including *C. dubliniensis* and *C. tropicalis* (data not shown).

**Figure 3.**
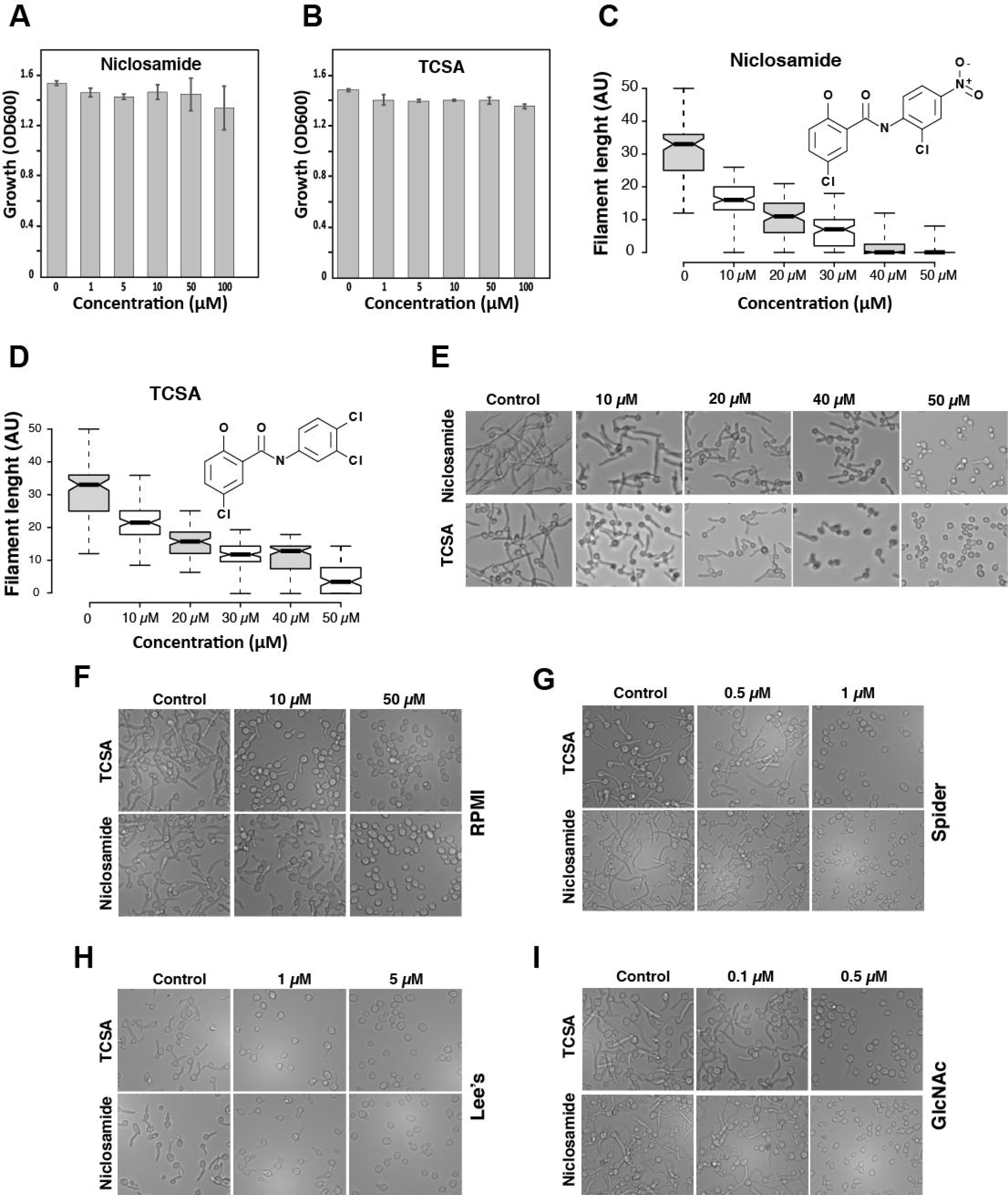
Niclosamide and TCSA abolished *C. albicans* hyphal growth promoted by different cues. Effect of Niclosamide (**A**) and TCSA (**B**) on the growth of the commensal yeast form. *C. albicans* SC5315 strain was exposed at different concentrations of the two HSA in SC medium and grown at 30°C for 24h. (**C**) Dose-response effect of Niclosamide (**C**) and TCSA (**D**) on filament length of *C. albicans* visualized by box plots. Bold lines in the box plots show the medians. *C. albicans* SC5315 strain was grown under hyphae-promoting conditions (SC + 10% FBS) in the absence (control) or the presence of HSA (10-50µM). Filament length were measured for at least 100 cells and shown as arbitrary unit (AU). (**E-I**) DIC micrographs showing the dose-response antifilamentation effect of Niclosamide and TCSA on *C. albicans* hypha grown on SC medium supplemented with 10% FBS serum (**E**) or on RPMI (**F**), Spider (**G**) and Lee’s (**H**) media, and SC with N-acetyl-D-glucosamine (GlcNAc) (**I**).

### TCSA and Niclosamide inhibit *C. albicans* biofilm formation

The effect of the two HSA were also assessed on *C. albicans* biofilm, a highly resistant sessile growth lifestyle associated with the contamination of medical devices. Niclosamide and TCSA impaired considerably the ability of *C. albicans* to establish biofilms (Fig. 4A-B). Antibiofilm activity were noticed at 1µM (12% inhibition) and 5µM (15% inhibition) for TCSA and Niclosamide, respectively. Both compounds were active on the biofilm of the emerging multi-drug resistant yeast, *C. auris*, and their inhibitory effect was perceived at 1µM (Fig. 4C-D). For both *Candida* species, biofilm inhibition was greater with TCSA as compared to Niclosamide.

**Figure 4.**
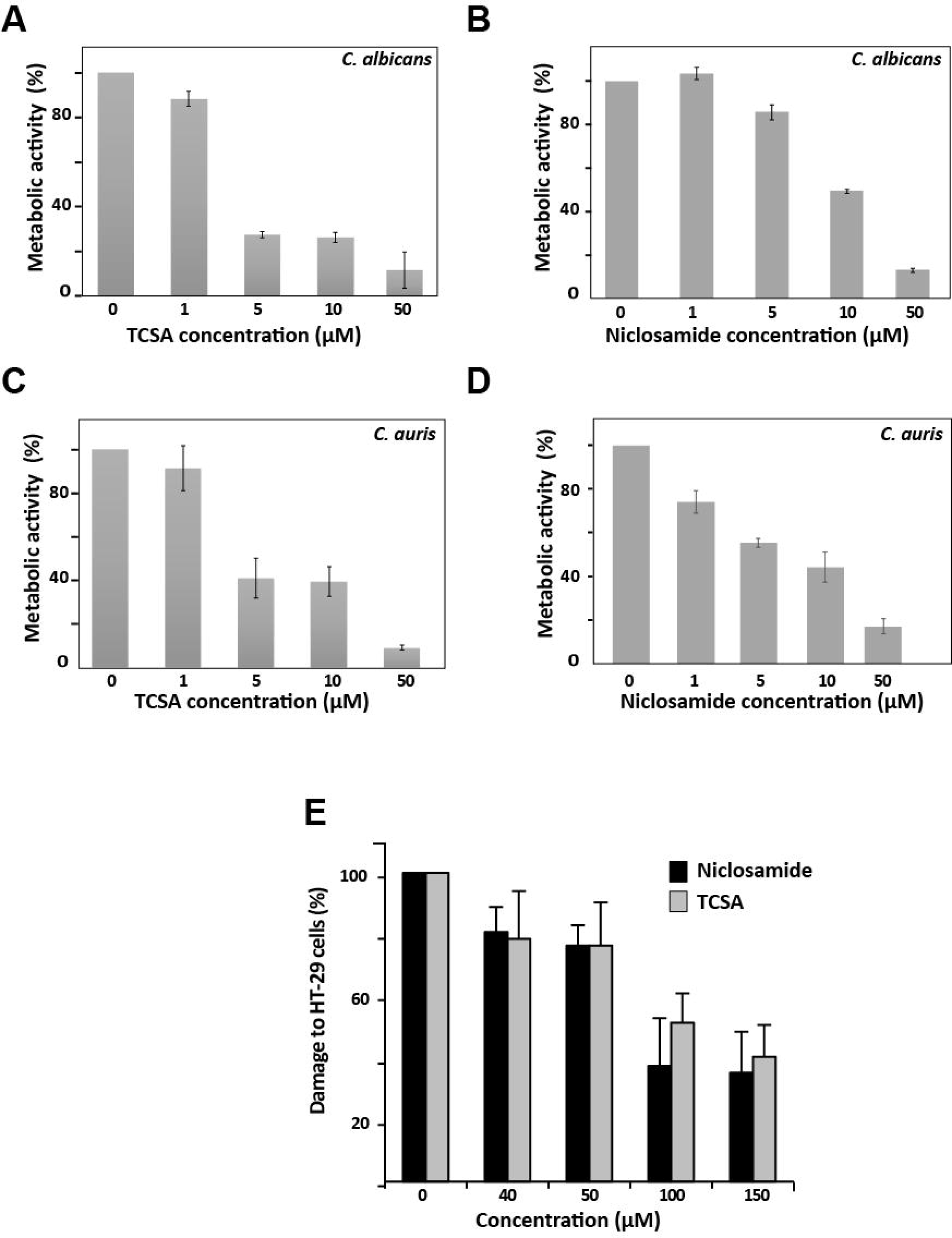
Niclosamide and TCSA antivirulence activity are expanded to the inhibition of biofilm formation and host invasion. Effect of TCSA (**A** and **C**) and Niclosamide (**B** and **D**) on biofilm formation of *C. albicans* (**AB**) and *C. auris* (**C-D**). The effect of HSA on biofilm formation of both *C. albicans* SC5314 and *C. auris* HDQ-RPCau1 strains was assessed using the metabolic colorimetric assay based on the reduction of XTT. Results represent growth inhibition (%) and are shown as the mean of at least three independent replicates. (**E**) Both Niclosamide and TCSA attenuate damage of enterocytes cells caused by *C. albicans*. Damage of the human epithelial intestinal cells HT-29 infected by *C. albicans* SC5314 strain was assessed using LDH release assay. Cell damage was calculated as percentage of LDH activity of HSA-treated experiment relatively to that of the control experiment (*C. albicans* invading HT-29 cells in the absence of HSA). Results are represented as the mean of three independent replicates.

### Niclosamide and TCSA attenuate damage of intestinal epithelial cells caused by *C. albicans*

Since Niclosamide and TCSA inhibited the formation of invasive hyphae, we wanted to check whether it conferred a protective activity for host cells against fungal invasion. *C. albicans-*mediated damage of the human colon epithelial HT-29 cells was quantified based on the LDH release assay in cells treated or not with different concentrations of the two HSA. Our data showed clearly that both antifilamentation compounds significantly reduced, but not completely abolished, the damage to HT-29 enterocytes (Fig. 4E). The protective effect was perceived at 40µM of either Niclosamide or TCSA (~ 20% reduction of enterocyte invasion). At concentration ≥100µM, Niclosamide and TCSA attenuated HT-29 cell damage by 63% and 49%, respectively.

### Niclosamide induces retrograde response in *C. albicans*

To gain insight into the mechanism of action associated with the antifilamentation effect of HSA, transcriptional profiling was undertaken on cells grown under hyphae-stimulating conditions in the presence of 50µM of Niclosamide. Gene ontology analysis showed that transcripts related to drug transport and, carbohydrate, trehalose and glyoxylate metabolisms were induced while genes of cell wall, filamentation and lipid metabolism including ergosterol (*ERG6*, *ERG25*, *CYB5, NDT80*), sphingolipids (*LCB2*, *FEN1*, *LAC1*) and phosphoglycerides (*AYR1*, *CHO2*, *PEL1*, *EPT1*, *YEL1*, *DGK1*) were downregulated (Table 1). We also found that genes encoding different proteins that control the activity of different protein kinases such as those modulating cell cycle progression (*CLN3*, *CLG1*, *SOL1*, *CIP1*) were repressed in response to Niclosamide. Gene expression alteration of eight genes including four upregulated (*ICL1*, *MLS1*, *MDR1*, *GDH2*) and four downregulated (*ZRT2*, *ERG25*, *LAC1*, *CDC47*) transcripts was confirmed using qPCR (**Table S1**).

**Table 1.**
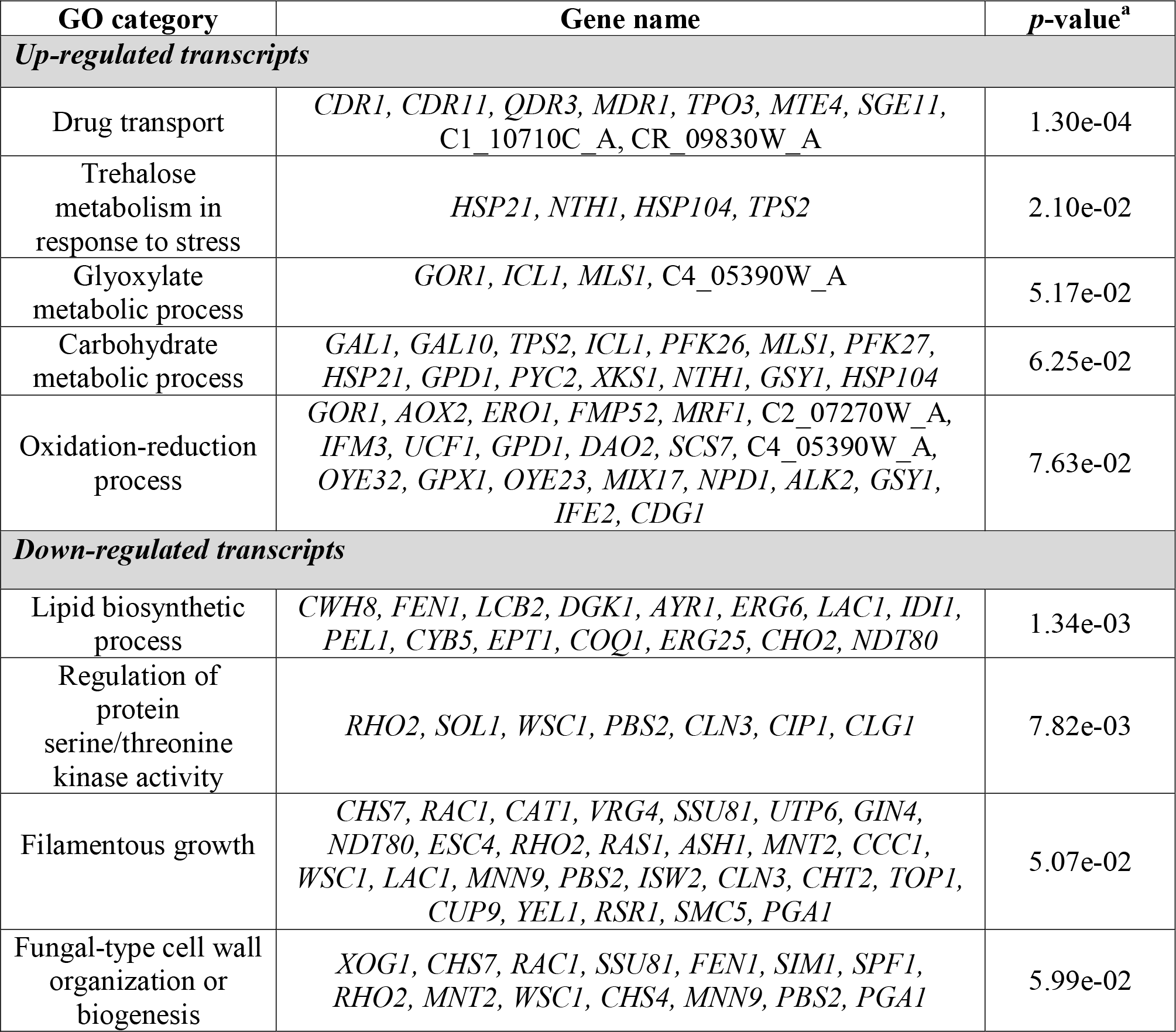
Gene function and biological process associated with *C. albicans* response to Niclosamide. Gene ontology analysis was performed using the Candida Genome Database GO Term Finder. The *p*-value was calculated using hypergeometric distribution, as described on the GO Term Finder website.

| GO category | Gene name | *p*-value<sup>a</sup> |
| --- | --- | --- |
| ***Up-regulated transcripts*** |  |  |
| Drug transport | *CDR1, CDR11, QDR3, MDR1, TPO3, MTE4, SGE11, C1_10710C_A, CR_09830W_A* | 1.30e-04 |
| Trehalose metabolism in response to stress | *HSP21, NTH1, HSP104, TPS2* | 2.10e-02 |
| Glyoxylate metabolic process | *GOR1, ICL1, MLS1, C4_05390W_A* | 5.17e-02 |
| Carbohydrate metabolic process | *GAL1, GAL10, TPS2, ICL1, PFK26, MLS1, PFK27, HSP21, GPD1, PYC2, XKS1, NTH1, GSY1, HSP104* | 6.25e-02 |
| Oxidation-reduction process | *GOR1, AOX2, ERO1, FMP52, MRF1, C2_07270W_A, IFM3, UCF1, GPD1, DAO2, SCS7, C4_05390W_A, OYE32, GPX1, OYE23, MIX17, NPD1, ALK2, GSY1, IFE2, CDG1* | 7.63e-02 |
| ***Down-regulated transcripts*** |  |  |
| Lipid biosynthetic process | *CWH8, FEN1, LCB2, DGK1, AYR1, ERG6, LAC1, IDI1, PEL1, CYB5, EPT1, COQ1, ERG25, CHO2, NDT80* | 1.34e-03 |
| Regulation of protein serine/threonine kinase activity | *RHO2, SOL1, WSC1, PBS2, CLN3, CIP1, CLG1* | 7.82e-03 |
| Filamentous growth | *CHS7, RAC1, CAT1, VRG4, SSU81, UTP6, GIN4, NDT80, ESC4, RHO2, RAS1, ASH1, MNT2, CCC1, WSC1, LAC1, MNN9, PBS2, ISW2, CLN3, CHT2, TOP1, CUP9, YEL1, RSR1, SMC5, PGA1* | 5.07e-02 |
| Fungal-type cell wall organization or biogenesis | *XOG1, CHS7, RAC1, SSU81, FEN1, SIM1, SPF1, RHO2, MNT2, WSC1, CHS4, MNN9, PBS2, PGA1* | 5.99e-02 |

Of note, transcripts of the isocitrate lyase, *ICL1*, and the malate synthase, *MLS1*, both key enzymes of glyoxylate cycle were highly induced (46- and 13-fold induction for *ICL1* and *MLS1*, respectively) (Fig. 5A and **Table S2**). The glyoxylate cycle is an anaplerotic pathway of the tricarboxylic acid (TCA) cycle that allows assimilation of carbon from C_2_ compounds by bypassing the CO_2_-generating steps of the TCA cycle. Both Icl1 and Mls1 enzymes are perxisomal, unique to the glyoxylate route and are not shared with the tricarboxylic acid (TCA) cycle ^35,36^. Activation of those genes in response to Niclosamide suggests a metabolic reconfiguration where acetyl-CoA is supplied by the glyoxylate cycle instead of the TCA cycle, which might be compromised (Fig. 5A). This transcriptional signature is characteristic of the retrograde (RTG) response, which is a mitochondria-to-nucleus signaling that cells activates as a cytoprotective mechanism to compensate for mitochondrial dysfunctions ^37^. Another supportive evidence of the TCA cycle failure, is reflected by the activation of genes that mediate other anaplerotic reactions to supply the TCA cycle with intermediates from alternative pathways. This includes the cytoplasmic pyruvate carboxylase enzyme, Pyc2, that converts the pyruvate to oxaloacetate. Transcript level of the glutamate dehydrogenase, Gdh2, that degrades glutamate to ammonia and alpha-ketoglutarate, were also induced (2.67-fold change), however, the levels of induction were not consistent enough (*p* = 0.21) for it to pass the threshold of statistical significance.

**Figure 5.**
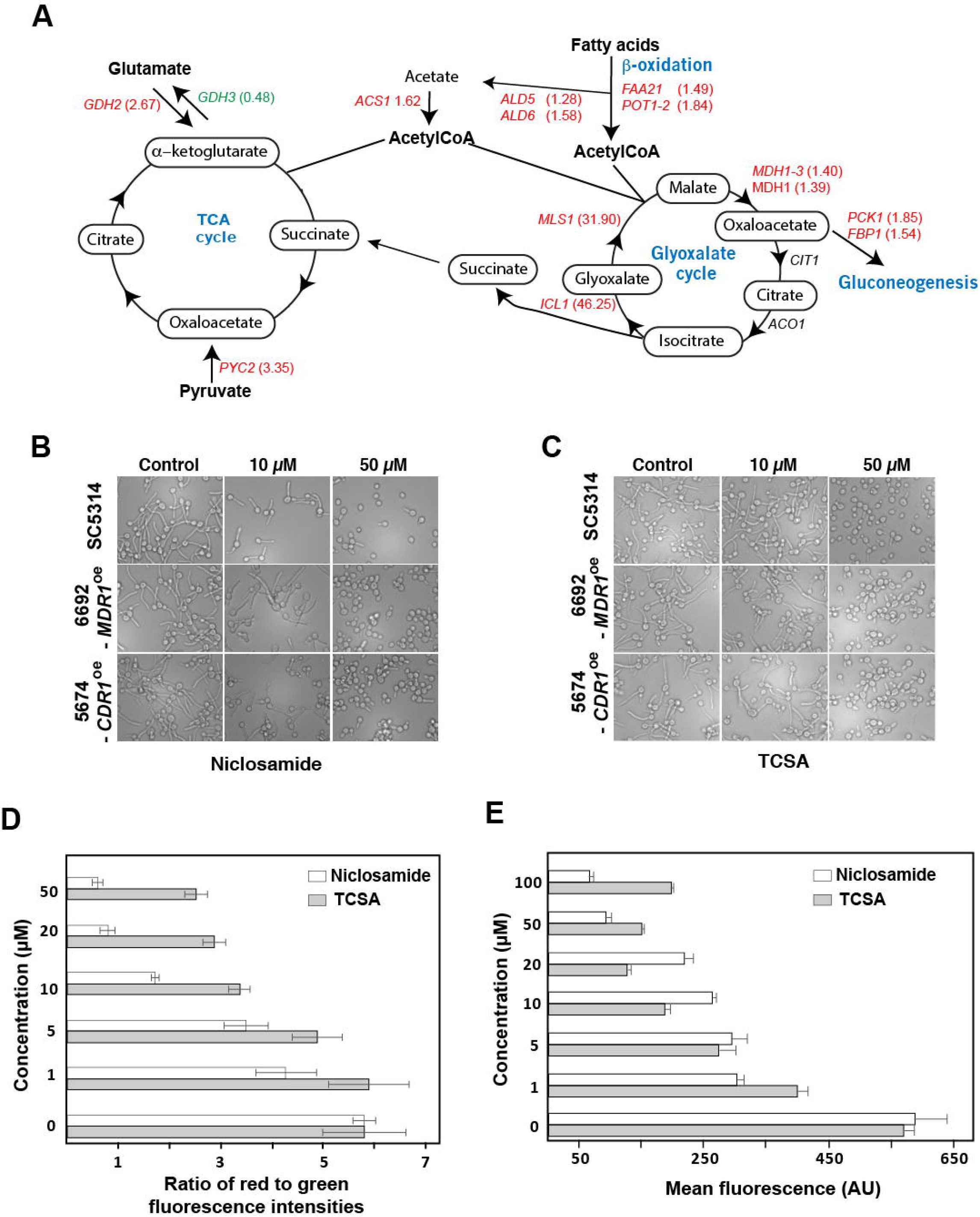
Niclosamide induces retrograde response in *C. albicans*. (**A**) Genome-wide transcriptional profiling reveals that transcript level of genes of anaplerotic reactions are altered in response to Niclosamide. Upregulated and downregulated genes are indicated by red and green, respectively. Transcripts that were not differentially expressed are shown in black. Simplified tricarboxylic and glyoxylate cycles as annotated in the CGD database are shown. (**B-C**) Antifilamentation effect of Niclosamide (**B**) and TCSA (**C**) on *C. albicans* resistant strains with different resistance mechanisms. *C. albicans* 6692 and 5674 resistant strains were grown under hyphae-promoting conditions (SC + 10% FBS) in the absence (control) or the presence of HSA (10-50µM). (**D-E**) Niclosamide and TCSA alter the mitochondrial membrane potential. *C. albicans* SC5314 strain cells were treated with different concentrations (1-50µM) of either TCSA or Niclosamide. Mitochondrial membrane potential (ΔΨm) were measured using the fluorescent potentiometric dye JC-1 and flow cytometry (**D**). Alternatively, the effect of both HSA on ΔΨm was assessed by MitoTracker staining (**E**). For both of JC-1 and MitoTracker assays, fluorescence ratio and intensities (AU) were presented as the mean of at least three independent experiments.

Activated transcripts were also enriched in genes encoding many proteins with oxidoreductase activity which might reflect a compensatory response to reoxidize the NADH as a consequence of TCA and mitochondrial electron transport failures. Reactivation of glycolytic regulatory genes *PFK26* and *PFK27* suggests also an adaptation to the drop of the intracellular level of ATP as a result of mitochondrial dysfunction. The strong upregulation of the alternative oxidase transcript, Aox2 (31.5-fold induction), emphasizes that *C. albicans* cells might use this alternative electron transport systems to bypass the lack of mitochondrial electron transport. Taken together, the Niclosamide-transcriptional signature is reminiscent of cells experiencing an RTG response as a consequence of impaired mitochondrial activity.

### Antifilamentation activity of Niclosamide and TCSA against *C. albicans* azole-resistant strains

In response to Niclosamide, *C. albicans* cells induced the expression of transporters such as Cdr1 and Mdr1 that are a key determinant of antifungal clinical resistance in this human pathogen ^38-40^. This reflect that *C. albicans* is trying to expel Niclosamide through efflux process. This prompt us to check whether antifungal resistance mediated by efflux process might bypass the antifilamentation effect of HSA. Both Niclosamide and TCSA were tested on clinical strains for which azole resistance is mediated by *MDR1* (strain 6692) and *CDR1* (strain 5674) overexpression. As shown in Fig. 5B-C, the antifilamentation effect of the two salicylanilides on these two resistant isolates was similar to that noticed for the SC5314 sensitive clinical strain. This emphasizes that Niclosamide and TCSA could be used to compromise *C. albicans* filamentation of both azole-sensitive and resistant strains.

### Niclosamide and TCSA alter the mitochondrial membrane potential

In the budding yeast, RTG response is triggered by the loss of mitochondrial membrane potential (Δψm) as a consequence of mitochondria dysfunction ^41^. Thus, Niclosamide or TCSA might induce RTG in *C. albicans* by promoting the perturbation of the Δψm. Change in Δψm was quantitatively assessed using JC-1, a green fluorescent potentiometric dye that shifts to red fluorescence as a consequence of Δψm. *C. albicans* cells treated with either Niclosamide or TCSA exhibited a collapse of the Δψm which was obvious at the concentration of 5µM (Fig. 5D). The effect of both HSA on the failure of Δψm was also confirmed using the MitoTracker dye that stains and accumulate in mitochondria dependently on Δψm ^42^ (Fig. 5E). These data suggest that both HSA induce RTG response in *C. albicans* as consequence of the alteration of Δψm.

### Mutants of the mitochondrial protein import machinery are required for Niclosamide tolerance and filamentation

Chemical-genetic assays such as the haploinsufficient profiling (HIP) is a powerful tool that has been widely used to uncover the mechanism of action (MoA) of many bioactive compounds ^43-46^. The principle of the HIP assay is based on the fact that decreased dosage of a drug target gene in a heterozygous mutant can result in increased drug sensitivity. To investigate the MoA of the HSA associated with their antifilamentation activity, we mined the data of HIP experiments performed in the yeast model *S. cerevisiae* in two resource studies ^47,48^. Hoepfner *et al*. ^47^ has performed HIP assay for Niclosamide while other analogs similar to HSA were investigated using the same approach by Corey Nislow’ group ^48^. To gain a consensual knowledge into the MoA of HSA, common hits identified by HIP assay of these related compound were identified by Venn diagram that revealed *mge1*/*MGE1* mutant as unique common hit for the five HSA (Fig. 6A).

**Figure 6.**
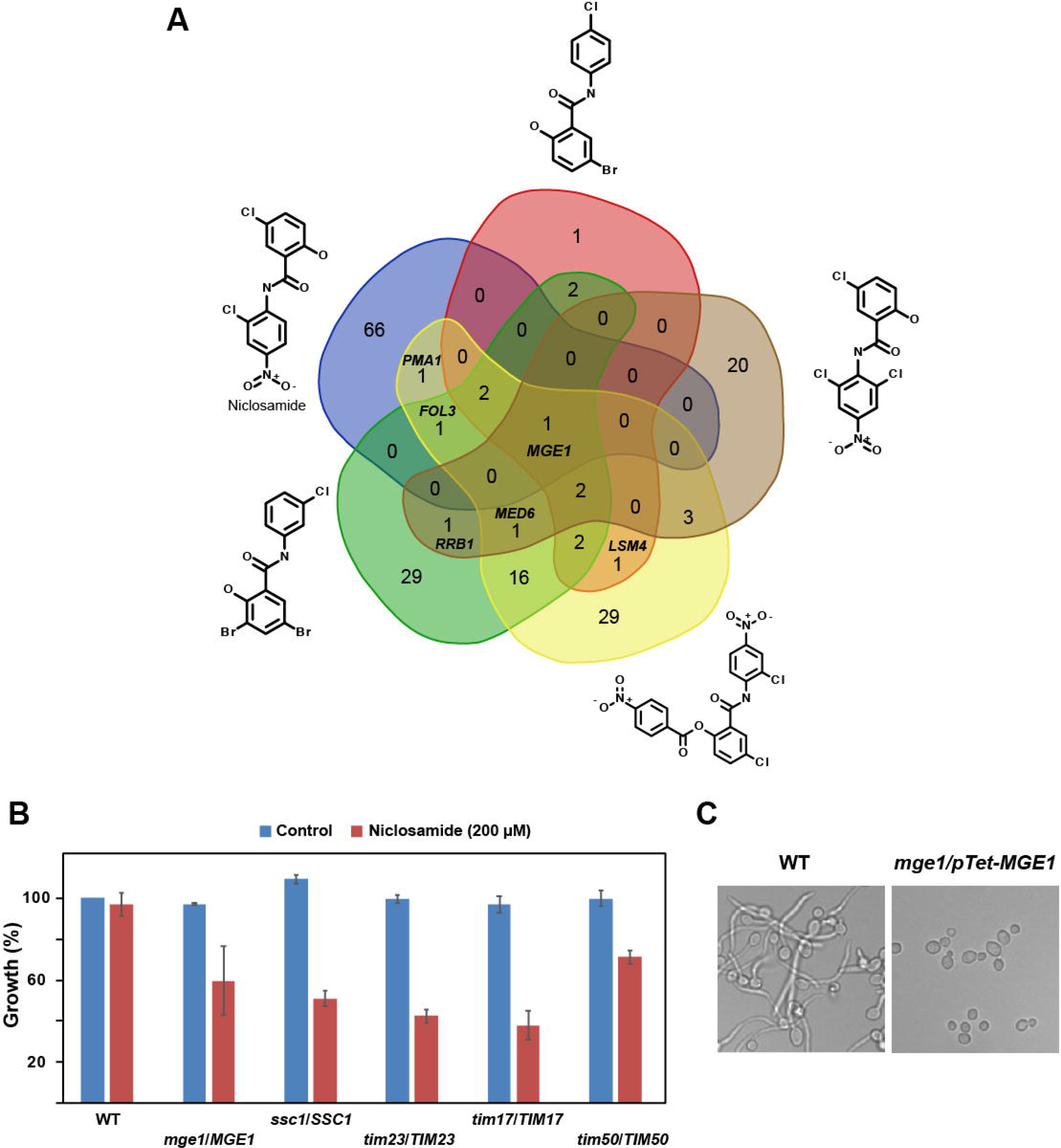
Mitochondrial protein import machinery is required for Niclosamide tolerance and filamentation. **(A)** Mining HIP assay data of HSA from studies performed by Hoepfner *et al*. ^47^ and Corey Nislow’ group ^48^ (http://chemogenomics.pharmacy.ubc.ca/hiphop/) in *S. cerevisiae*. HSA-haploinsufficient mutant hits identified by HIP assay for each compound were selected based on *z*-score ≥ 2.5. The Venn diagram was generated using the web tool software at the following URL: www.bioinformatics.psb.ugent.be/webtools/Venn. **(B)** *C. albicans* heterozygous mutants of the TIM23 complex including *tim23*/*TIM23*, *tim17*/*TIM17, tim50*/*TIM50, ssc1*/*SSC1* and *mge1*/*MGE1* are sensitive to high concentration of Niclosamide (200µM). **(C)** The conditional homozygous mutant *mge1/pTeT-MGE1* under repressive conditions (100µg/mL tetracycline) is unable to differentiate true hyphae when grown at 37°C in Spider medium as compared to the parental strain CAI4.

Mge1 is a nucleotide release factor for the mitochondrial heat shock protein 70, Ssc1 that acts by facilitating the release of ADP from Ssc1 ^49^. This process is essential for the import complex to translocate nuclear-encoded proteins to their final mitochondrial destination. Both Mge1 and Scc1 are components of the Tim23-Tim17 import motor complex that mediates the translocation of proteins through the inner mitochondrial membrane ^50^. The sensitivity of *C. albicans* heterozygous mutants of TIM23 complex including *tim23*/*TIM23*, *tim17*/*TIM17, tim50*/*TIM50, ssc1*/*SSC1* and *mge1*/*MGE1* was tested toward high concentration of Niclosamide (200µM). While the WT parental strain had no discernable growth defect, all tested mutants were sensitive when exposed to Niclosamide (Fig. 6B). The Niclosamide-induced haploinsufficiency of the TIM23 complex suggests that the *C. albicans* translocase machinery of the inner mitochondrial membrane is required to tolerate Niclosamide and/or might be a target of this HSA.

In opposite to *S. cerevisiae, MGE1* is not essential in *C. albicans* ^51^. As for the heterozygous mutant strain, the conditional homozygous mutant *mge1/pTeT-MGE1* under repressive condition (100µg/mL tetracycline) was also sensitive to Niclosamide (not shown). Furthermore, genetic inactivation of *MGE1* led to a complete filamentation defect which support the hypothesis that this protein might be the target of HSA (Fig. 6C).

### Chemical genetics of hyphal signaling using TCSA and Niclosamide as perturbers

The *C. albicans* morphogenetic switch is controlled by intertwined regulatory circuits that signal different filamentation cues. The RAS/cAMP ^52^, the MAPK ^53^ and Ume6/Hgc1 ^54^ pathways are high hierarchical regulators and signaling hubs that control the filamentation process in *C. albicans*. To test whether Niclosamide and TCSA alter *C. albicans* filamentation through these pathways, the effect of the two molecules was assessed in mutants overexpressing *UME6*, *HGC1* and the MAPKKK, *STE11* in addition to the dominant mutant Ras1^G13V^. These mutants constitutively formed hyphae or pseudohyphae which will determine whether antifilamentation molecules act upstream or downstream these pathways. Niclosamide and TCSA blocked *C. albicans* filamentation regardless the targeted regulators suggesting that both compounds acted on effectors downstream these signaling pathways (Fig. 7A-B).

**Figure 7.**
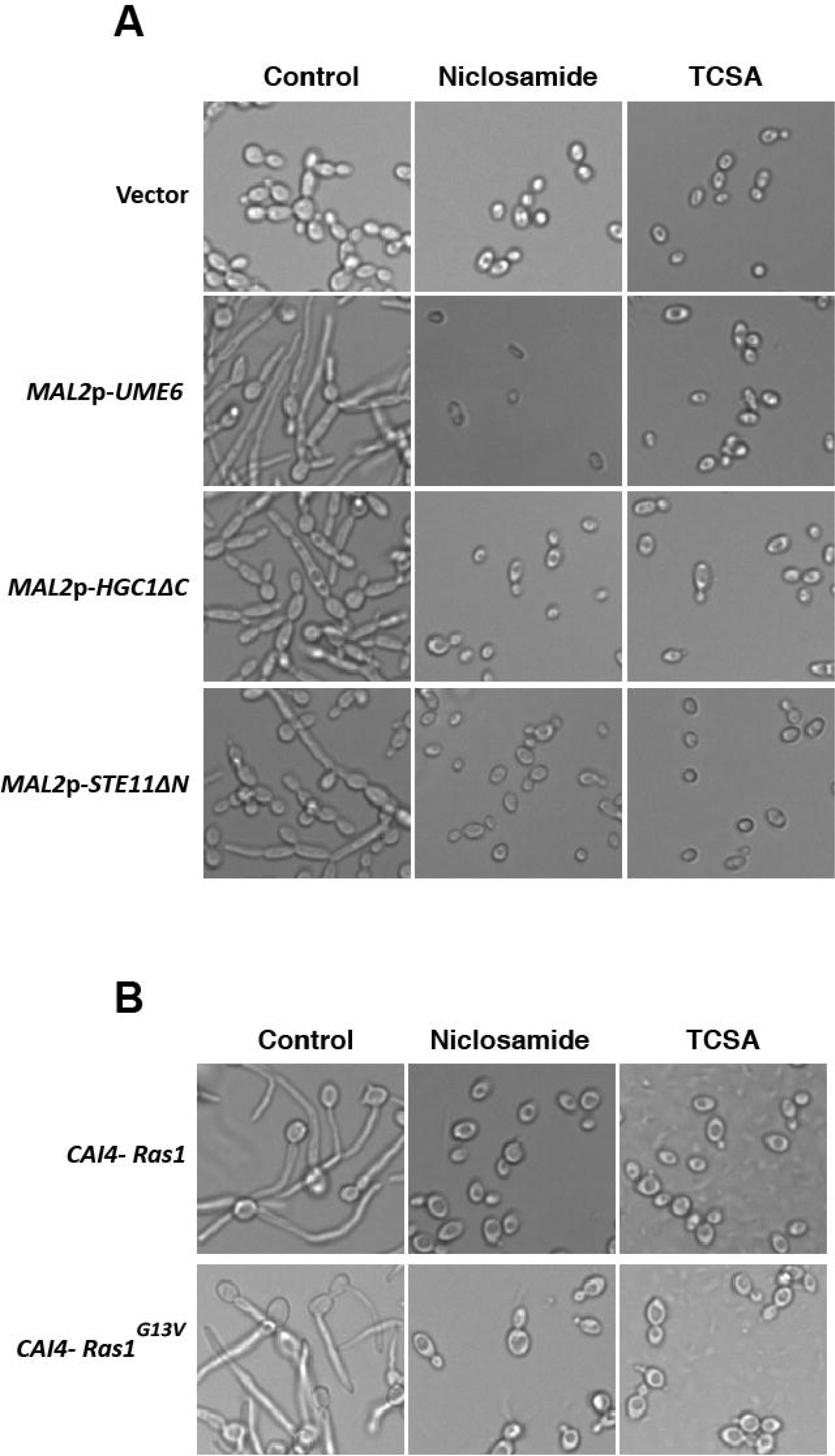
Chemical genetics of the *C. albicans* hyphal signaling network using HSA as perturbers. (**A**) Effect of Niclosamide and TCSA on *C. albicans* mutants overexpressing key regulator of the yeast-to-hyphae transition. Ectopic filamentation of *MAL2p-UME6*, *MAL2p-HGC1*Δ*C* and *MAL2p-STE11ΔN* mutants was ensured by growing these strains under inducing conditions in YNB-2% maltose medium at 30°C for 3 hours. (**B**) Constitutive filamentation of the dominant Ras1^G13V^ mutant is inhibited by Niclosamide and TCSA. Both Ras1^G13V^ mutant and the control strain (CAI4-Ras1) were grown at 30°C on Spider medium.

## Discussion

Neutralizing the virulence traits of a fungal pathogen represents a promising therapeutic strategy to circumvent antifungal resistance. Regardless the degree of susceptibility or resistance of a fungal strain toward conventional antifungal drugs, antivirulence molecules lead to the silencing of the virulence machinery which in turn inhibits the installation of the infection. Furthermore, antivirulence agents could bypass the undesirable side effects of antimicrobials that kill beneficial commensal fungi including *C. albicans* itself ^13^. In the current work, we have repurposed the widely used anthelminthic, Niclosamide and its derivative TCSA as a promising antivirulence compounds that impede both invasive filamentation and biofilm formation in both susceptible and resistant clinical *C. albicans* strains. Currently, the antivirulence paradigm was clinically exploited to manage bacterial virulence factors such as anthrax toxins and botulinum neurotoxin ^13^. To our knowledge, and given the fact that the safety of Niclosamide is well established in humans ^55^, this molecule could represent the first clinically approved antivirulence agent against a pathogenic fungus. Interestingly, this anthelmintic agent was also shown to be affective toward bacterial pathogens as an antivirulence molecules against *Pseudomonas aeruginosa* ^56^ or as an antibiotic against multi-resistant strains of *Staphylococcus aureus* ^26^. This newly discovered antibacterial and antifungal proprietes would reinforce and maintain the status of Niclosamide as being one of the essential medicines in the list the World Health Organization.

Here, we uncovered that halogenated salicylanilides represent new chemical scaffolds that impede specifically *C. albicans* virulence without affecting the commensal yeast growth. Future comprehensive structure-guided medicinal chemistry investigations are required to precisely point out the chemical group that lead to this discriminative antivirulence activity. Of note, the non-halogenated salicylanilide scaffold (compound #3) has a strong anti-growth activity against *C. albicans* and suggests that halogenation enhances the antivirulence activity. Halogen bonds are exploited in medicinal chemistry for their steric effect to accommodate a best fitting of a small molecule to occupy the binding site of its target ^57^. Our data showed that chlorination was more effective as compared to fluorination (compound #5 and #8) or bromination (compound #11) regarding the antivirulence activity. This might be explained by the fact that, as compared to the other halogens-carbon bonds, C-Cl is more stable which might allow a steady docking on its virulence-related target ^57^.

Regarding the mechanism of action, HSA have been associated with the perturbation of a plethora of biological processes in bacteria and parasites. Salicylanilides have been shown to inhibit the two-component regulatory systems, uptake of nutrients and oxidative phosphorylation, and to cause leakage and membrane damage and protein aggregation ^58^. However, their mechanisms associated with their antifungal or antivirulence activity were not investigated so far. In the current study, the transcriptional signature exhibited by *C. albicans* cells exposed to Niclosamide suggests activation of different anaplerotic metabolic pathways that can sustain the TCA cycle with metabolites such as citrate, succinate and oxaloacetate. This phenomenon, so-called RTG, is most likely triggered in *C. albicans* cells as a consequence of the dissipation of the mitochondrial potential. The HIP assays on cells treated to different HSA uncovered that Mge1 might be a potential target of Niclosamide and TCSA. Indeed, genetic inactivation of Mge1 led to a complete filamentation defect which supports the hypothesis that this protein might be the target of HSA.

Mge1 is a nucleotide release factor that mediates the release of ADP from the heat shock protein Ssc1 ^49^. This process is essential for the mitochondrial import complex to translocate nuclear-encoded proteins to mitochondria. Interestingly, this process is Δψm-depend which suggests that the collapse of Δψm mediated by HSA might contribute to the malfunctioning of the mitochondrial import complex and thus compromises protein translocation in the mitochondria ^59^. Alternatively, in addition to its role as ADP-ATP recycling factor, Mge1 functions as a co-chaperone in the mitochondrial matrix together with Scc1 and Mdj1 to refold desaturated proteins ^60^. Taken together, either protein translocation or refolding functions of Mge1 could be hampered by HSA which in turn lead to the filamentation defect in *C. albicans*.

While deep investigations are required to validate that Mge1 and/or the mitochondrial protein import complex as a target of HSA, previous elegant investigations in *C. albicans* have shown that many fungal mitochondrial proteins are therapeutic targets for antifungal development ^61^. Furthermore, Mge1 has been recently shown to be a key regulator of azole susceptibility ^62^ which make the fungal mitochondria as an attractive organelle target to develop anti-virulence, anti-growth and anti-resistance molecules to manage fungal infection.

## Methods

### Strains and growth conditions

The strains used in the current study are listed and described in Table 2. For general propagation, the strains were cultured at 30°C in synthetic complete (SC; 0.67% yeast nitrogen base with ammonium sulfate, 2% glucose, and 0.079% complete supplement mixture) or yeast-peptone-dextrose (YPD; 2% Bacto-peptone, 1% yeast extract, 2% dextrose) media supplemented with uridine (50mg/L).

**Table 2.** Fungal strains used in this study.

| Name | Description/Genotype | Reference |
| --- | --- | --- |
| ***Candida albicans*** |  |  |
| SC5314<br>(ATCC-MYA-2876) | *C. albicans* wild-type reference strain. | 70 |
| 5674 | Azole-resistant clinical strain (overexpressing the ABC-transporters Cdr1 and Cdr2, and the phosphatidylinositol transfer protein, Pdr16) isolated from mouth | 71 |
| 6692 | Azole-resistant clinical strain (overexpressing the MFS-transporter Mdr1 and had a gain-of-function mutation on the transcription factor, Mrr1) isolated from mouth | 71,72 |
| KC685 | *ura3Δ/ura3Δ his1Δ/his1Δ leu2Δ/leu2Δ arg4Δ/arg4Δ pMAL2-URA3* | 16 |
| KC753 | *KC685 STE11/pMAL2- STE11ΔN-URA3* | 16 |
| KC763 | *KC685 UME6/pMAL2-UME6-URA3* | 16 |
| KC754 | *KC685 HGC1/pMAL2- HGC1ΔC-URA3* | 16 |
| DH545 | *ura3::λimm434/ura3 pYPB1-ADHpL-CaRAS1* | 73 |
| DH409 | *ura3::λimm434/ura3 pYPB1-ADHpL-CaRAS1<sup>G13V</sup>* | 73 |
| ***Candida auris*** |  |  |
| HDQ-RPCau1 | Clinical isolate from Hôtel-Dieu de Québec Hospital, QC, Canada | - |

### Screen of the bioactive chemical yeast library

To identify small molecules with antifilamentation properties, the yeast bioactive small molecule library from Dr Mike Tyers (University of Montreal) consisting of 678 compounds preselected from the Maybridge collection (Thermo Fisher Scientific) was screened. *C. albicans* SC5314 strain cells were grown in 96-well in SC medium supplemented with 10% FBS (Wisent) and one microliter of each compound from the master library plate was dispensed into the assay plate at a final concentration of 100µM. DMSO at 1% (v/v) final concentration was added as control to each assay plate. After an incubation for 3h in the high-content microscope Cytation 5 (BioTek^®^, Thermo Fisher Scientific), images were taken in each well. Hit from this primary screen were confirmed using new batch of powders ordered separately. Frequent hitters and promiscuous compounds in antifungal discovery pipelines ^21^ were removed from our hit list. Effect of antifilamentation compounds on the yeast growth of *C. albicans* were assessed as follow: SC5314 strain was grown overnight in YPD medium at 30°C in a shaking incubator at 220 rpm. Cells were then resuspended in fresh SC at an OD_600_ of 0.05. A total volume of 99μl fungal cells was added to each well of a flat-bottom 96-well plate in addition to 1μl of the corresponding stock solution antifilamentation molecules. Plates were incubated in a Sunrise-Tecan plate reader at 30°C with agitation and OD_600_ readings were taken every 10min over 24h. Each experiment was performed in triplicate.

### Hyphal growth and biofilm assays

An overnight culture of the *C. albicans* strains was used to inoculate a 5mL of fresh YPD at an OD_600_ of 0.05. Cells were grown for 4 h at 30°C under agitation to reach the exponential phase. To induce hypha, the SC5314, 6692 and 5674 strains were grown at 37°C in YPD supplemented with either 10% FBS (Wisent) or 2.5mM N-acetyl-D-glucosamine (Sigma), or in Spider (1% mannitol, 1% nutrient broth, 0.2% K2HPO4), Lee’s ^63^ or RPMI 1640 (Gibco) media. The filament length of the generated filaments was assessed for at least 100 *C. albicans* cells per sample, and all experiments were performed in triplicate. The effect of HSA on hyphae lengths were presented as box plot using the BoxPlotR web tool ^64^.

Biofilm formation and XTT (2,3-bis(2-methoxy-4-nitro-5-sulfo-phenyl)-2H-tetrazolium-5-carboxanilide) assays were carried out as follow: overnight YPD cultures were washed three times with PBS and resuspended in fresh RPMI 1640 supplemented with L-glutamine (0.3g/L) to an OD_600_ of 0.1. *C. albicans* yeast cells were allowed to attach to a flat-bottom 96-well polystyrene plate for 3h at 37°C in a rocking incubator. After removing non-attached cells by washing 3 time using PBS, fresh RPMI supplemented with HAS was added. The plates were then incubated for 24 hours at 37°C for biofilm formation under agitation at 220 rpm. Each well in the plate was then washed 3 times with PBS and fresh RPMI supplemented with 100μl of XTT-menadione mix (0.5mg/ml XTT in PBS and 1mM menadione in acetone) was added. After 3 hours of incubation on dark at 37°C, 80μl of the resulting colored supernatants were used for colorimetric reading at an OD_490_ to assess metabolic activity of biofilms. A minimum of four replicates were at least performed. Biofilm formation for *C. auris* was performed as for *C. albicans* with the sole exception that cells were allowed to attach for 6 hours.

### Transcriptional profiling

Overnight cultures of *C. albicans* strain SC5314 were diluted to an OD_600_ of 0.1 in 100mL of fresh YPD and grown at 30°C until an OD_600_ of 0.65. The culture was divided into two volumes of 50mL where 10% FBS serum was added to induce hyphae formation; one sample was maintained as the control where DMSO was added, and the other treated with Niclosamide at 50µM. *Candida* cells were then incubated at 37°C and under agitation at 220 rpm for 15 min. Cells were then centrifuged for 2min at 3,500 rpm, the supernatants were removed, and the samples were quick-frozen and stored at 80°C.

RNA extractions, cDNA labelling and microarrays procedures were performed as described by Chaillot *et al*. ^65^. Data analysis were carried out using Genespring v.7.3 (Agilent Technologies, Palo Alto, CA). Statistical analysis used Welch’s *t-*test with a false-discovery rate (FDR) of 5% and 2-fold enrichment cut-off. Gene ontology (GO) analysis was performed using the Candida Genome Database (CGD) GO Term Finder. The *p*-value was calculated using hypergeometric distribution with multiple hypothesis correction (http://www.candidagenome.org/cgi-bin/GO/goTermFinder) ^66^. Descriptions of each *C. albicans* gene in **Table S1** were extracted from CGD database ^67^.

For quantitative real time PCR (qPCR) confirmation, cell cultures and RNA extractions were performed as described for the microarray experiment. cDNA was synthesized from 1µg of total RNA using High-Capacity cDNA Reverse Transcription kit (Applied Biosystems). The mixture was incubated at 25°C for 10min, 37°C for 120min and 85°C for 5min. Then RNAse H (NEB) was added to remove RNA. qPCR was performed using LightCycler 480 Instrument (Roche Life Science) with SYBR Green fluorescence (Applied Biosystems). The reactions were incubated at 50°C for 2 min, 95°C for 2min and cycled 40 times at 95°C, 15 s; 54°C, 30 s; 72°C, 1 min. Fold-enrichment of each tested transcripts was estimated using the comparative ΔΔCt method as described by Guillemette *et al*. ^68^. To evaluate the gene expression level, the results were normalized using Ct values obtained from Actin (*ACT1*, C1_13700W_A). Primer sequences used for this analysis are summarized in Supplemental Table S3.

### HT-29 damage assay

Damage to the human enterocyte HT-29 (ATCC-HTB-38) was assessed using a lactate dehydrogenase (LDH) cytotoxicity detection kit, which measures the release of LDH in the growth medium as described by Garcia *et al*. ^69^. Briefly, enterocytes were plated at 10.000 cells per 96-well in Dulbecco’s modified Eagle’s medium (DMEM) supplemented with 10% FBS and incubated overnight at 37°C with 5% CO_2._ The HT-29 cells were then infected with *C. albicans* SC5314 strain cells with a multiplicity of infection (MOI) of 1:20 (Enterocyte:*C. albicans*). Following 24 hours of incubation, 100µL supernatant was removed from each experimental well and LDH activity in this supernatant was determined by measuring the absorbance at 490 nm. Each LDH activity assay was performed in triplicate and at least three independent biological replicates were performed.

### Chemical epistasis assay

Inhibition assay of ectopically hyphae-grown strains KC753 (*MAL2-*STE11Δ*N*), KC763 (*MAL2-UME6*) and KC754 (*MAL2-HGC1ΔC*) as well as their control strain KC685 was performed as follow: cells were grown overnight in SC medium where glucose was replaced by 2% raffinose. Overexpression of *UME6*, *STE11* and *HGC1* was ensured by diluting the overnight cultures in fresh SC medium where glucose was replaced by 2% maltose in the presence or not of HSA. The cells were incubated for 5h at 30°C. The DH409 mutant with a hyperactive Ras1 (Ras1^G13V^) allele and the control strain DH545 were allowed to filament in Spider medium at 30°C in the presence of HSA for 2h.

### Measurements of mitochondrial membrane potential

The mitochondrial membrane potential of *C. albicans* cells was measured using both fluorescent probes, JC-1 and MitoTracker Red CMXRos (ThermoFisher). Exponentially grown *C. albicans* cells were washed twice with PBS and treated or not with different concentrations of either Niclosamide or TCSA for 30 min under hyphae promoting conditions. JC1 and MitoTracker were added to *C. albicans* cells at final concentration of 5 µM and 100 nM, respectively, and cells were incubated at 30°C for 30 min in dark. Cells were then washed twice with PBS to remove residual dyes. Fore JC-1, fluorescence was quantitatively assessed using flow cytometry (BD FACSCanto™) using 10^6^ *C. albicans* cells. For the MitoTracker assay, the fluorescence was measured using the Cytation 5 plate reader (BioTek^®^). Results of JC-1 and MitoTracker assays were measured as the mean of fluorescence intensities of at least three independent experiments.

## Acknowledgments

We are grateful to Mike Tyers and Susan Moore (Université de Montreal) for providing the yeast bioactive small molecule library. We thank Martine Raymond (Université de Montreal) for providing *C. albicans* clinical strains, René Pelletier (Research Center of the CHU de Québec) for the *C. auris* strain, Deborah Hogan (Geisel School of Medicine at Dartmouth) for sharing the Ras1^G13V^ mutant and Daniel Kornitzer (Technion) for the hypofilamentous *C. albicans* strains.

Work in Sellam’s laboratory is supported by the Fonds de Recherche du Québec-Santé (FRQS) (Établissement de jeunes chercheurs), the Canada Foundation for Innovation (CFI-# 34171), the Fondation de l’Université Laval-Merck fund and the Natural Sciences and Engineering Research Council of Canada (NSERC) discovery grant (06625). AS is a recipient of the Fonds de Recherche du Québec-Santé (FRQS) J1 salary award. AB and JC received Ph.D. scholarships from Université Laval (Faculty of Medicine) and the CHUQ foundation.

## Author Contributions

AS designed the experiments. CG, AB, JC, EP, IK and AS performed the experiments and interpreted the data. CG and AS wrote the paper. All authors reviewed the manuscript.

